# Impact of Equilibrative Nucleoside Transporters on *Toxoplasma gondii* Infection and Differentiation

**DOI:** 10.1101/2024.07.10.601519

**Authors:** Gabriel Messina, Amber Goerner, Charlotte Bennett, Euwen Brennan, Vern B. Carruthers, Bruno Martorelli Di Genova

**Affiliations:** Department of Microbiology and Molecular Genetics, University of Vermont, Burlington, VT 05405, USA; Department of Microbiology and Immunology, University of Michigan School of Medicine, Ann Arbor, MI 48109, USA

## Abstract

*Toxoplasma gondii* cannot synthesize purines de novo and must import them; yet, the functional interplay among its four equilibrative nucleoside transporters (ENTs) homologs remains unclear. We systematically deconstructed this network by combining CRISPR-Cas9 knockouts with an auxin-inducible degron. Across all phenotypic assays, tachyzoite replication, nucleoside-analogue sensitivity, alkaline-stress–induced differentiation, and murine cyst formation, the Δ*TgENT2* strain was indistinguishable from the parental line, indicating that TgENT2 is dispensable under the conditions tested. In contrast, the double mutant Δ*TgAT1ΔTgENT3* exhibited delayed bradyzoite differentiation in vitro and produced smaller brain cysts in vivo. This double deletion triggered a ∼3-fold transcriptional up-regulation of *TgENT1*, whose product we partially localized to the plant-like vacuolar compartment (PLVAC). Conditional depletion of TgENT1 caused complete intracellular growth arrest, PLVAC swelling, and a purine-starvation-like transcriptomic program enriched for nucleoside phosphatases and cyclic-nucleotide phosphodiesterases. These findings reveal a compensatory salvage pathway in which the parasite reroutes purine acquisition through a vacuolar route when plasma-membrane import is compromised. Although this adaptation sustains tachyzoite proliferation, it fails during the energetically demanding transition to bradyzoites, creating a metabolic bottleneck that impairs chronic infection. Our work reveals an adaptable yet ultimately limited purine-import network and identifies TgENT1, along with the vacuolar salvage axis it mediates, as a promising therapeutic target for blocking lifelong toxoplasmosis.

## Introduction

*Toxoplasma gondii* is an obligate intracellular parasite that causes toxoplasmosis, a disease affecting approximately one-third of the global population and posing risks to immunocompromised individuals and developing fetuses. The parasite’s lifecycle includes acute (tachyzoite) and chronic (bradyzoite) stages, each characterized by distinct metabolic demands ^1^. A critical adaptation of *T. gondii* is its complete reliance on host-derived purines due to the absence of a *de novo* purine synthesis pathway ^2^. Understanding how *T. gondii* transports and manages purine resources is crucial for deciphering its survival strategies and identifying potential therapeutic targets. Equilibrative nucleoside transporters (ENTs) are key proteins involved in this purine acquisition from host cells ^3^. Here we investigate the roles of four putative *T. gondii* ENTs: TgENT1 (TGME49_288540), TgENT2 (TGME49_500147), TgENT3 (TGME49_233130), and TgAT1 (TGME49_244440) across both acute and chronic stages, aiming to elucidate their distinct and overlapping functions in parasite survival and development.

*De novo* nucleic acid synthesis and salvage pathways are fundamental for cellular growth and survival, allowing cells to adapt to varying environmental conditions ^4^. Higher eukaryotes possess sophisticated mechanisms to modulate nucleoside transport, salvage, and metabolism, integrating intracellular and extracellular cues to maintain nucleotide homeostasis ^5–9^.

Many early branching eukaryotes have evolved complex mechanisms to adapt to fluctuating nucleoside availability ^10–12^. *T. gondii* exemplifies this adaptation through its complete dependence on host-derived purines, having lost the capacity for *de novo* purine synthesis during evolution ^2,11,13^. This reliance underscores the importance of understanding the mechanisms of purine acquisition in *T. gondii*. In host cells such as macrophages and neurons, purine levels are tightly regulated, presenting a challenging environment for the parasite ^9,14^.

The parasite employs equilibrative nucleoside transporters (TgENTs) as its primary means of scavenging purines from the host ^15^. *T. gondii* utilizes two purine salvage pathways: the phosphorylation of adenosine by the enzyme adenosine kinase and the phosphoribosylation of purine nucleobases by hypoxanthine-guanine phosphoribosyltransferase (HXGPRT) ^11^. These pathways enable the parasite to salvage both adenosine and other purine nucleobases efficiently. Additionally, *T. gondii* possesses a functional *de novo* pyrimidine biosynthesis pathway, with uracil phosphoribosyltransferase (UPRT) facilitating pyrimidine salvage by converting uracil to uridine monophosphate ^11^.

The initial identification of equilibrative nucleoside transporter (ENT) activity in *T. gondii* was demonstrated in extracellular tachyzoites, where adenosine, inosine, and hypoxanthine were transported into the parasite ^16^. Subsequent studies identified TgAT1 via insertional mutagenesis^17^. Radioactive nucleoside incorporation assays revealed the presence of multiple nucleoside transporters in *T. gondii*, suggesting a complex system with at least one high-affinity adenosine transporter yet to be characterized ^18^. TgAT1 remains the only TgENT studied *in vitro* using heterologous expression systems ^15^.

In this study, we aim to elucidate the roles of TgAT1 and three other homologs, TgENT1, TgENT2, and TgENT3, across different developmental stages of *T. gondii*. We hypothesized that the network of purine transporters in *T. gondii* is characterized by functional redundancy and compensatory regulation, ensuring parasite survival under purine stress. To test this, we employed a systematic genetic approach to deconstruct this network by the investigation of the parasite’s adaptive response to each TgENT disruption; and assess the importance of this transporter network for the critical developmental transition from the acute tachyzoite to the chronic bradyzoite stage. This strategy was designed to unmask the latent functions and regulatory connections within the purine acquisition machinery.

## Results

### TgENT BLAST Analysis

We performed protein-protein BLAST or position-specific iterated BLAST on NCBI using Human ENT-1^3^ and *Plasmodium falciparum* PfENT1^12^ nucleoside transporters as query sequences to search for *T. gondii* sequences with high similarity. Phylogenetic analysis (**Fig. 1A**) places TgAT1 and TgENT1–3 in a single, well-supported clade with apicomplexan and mammalian ENTs. Conserved-domain searches and AlphaFold modelling suggest that TgAT1 and TgENT1-3 harbor the hallmark 11-ENT fold with an open central pore (**Fig. 1B,C**, and **Table 1**). Pairwise structural comparisons to the HsENT1 crystal structure (PDB 6OB7) show pore dimensions within the expected range for functional ENT homologues (**Fig. 1C**), supporting their candidacy as bona fide transporters. Finally, re-analysis of the transcriptomic data^19,20^ (**Fig. 1D**) reveals distinct but overlapping expression patterns; TgAT1 and TgENT1 are abundant throughout enteric, tachyzoite, and bradyzoite stages, TgENT3 is moderately expressed across the life cycle, and TgENT2 peaks during sporulation, suggesting stage-specific expression. A multiple-sequence alignment of TgENT1, TgENT2, TgENT3, TgAT1, and other characterized ENT homologs from different organisms is provided in Supplementary **Figures S1**.

**Figure 1.**
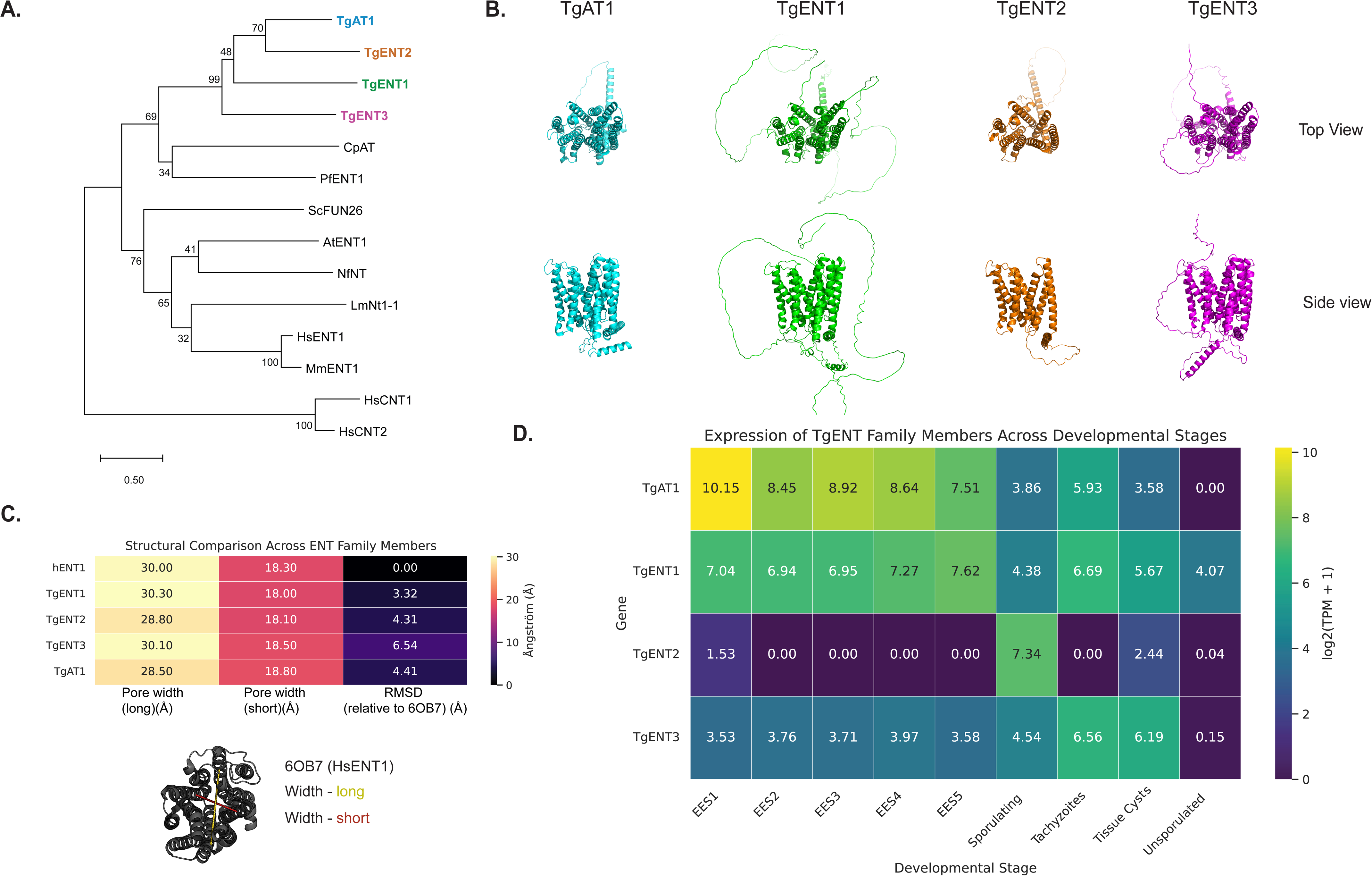
Evolutionary, structural, and transcriptional overview of the *Toxoplasma gondii* equilibrative nucleoside transporter (TgENT) family. **(A)** Phylogenetic relationships: Full-length protein sequences of the four *T. gondii* ENTs (TgAT1, TgENT1–3) were aligned with representative ENT homologs from apicomplexan, fungal, plant, trypanosomatid, and mammalian lineages (CpAT, *Cryptosporidium parvum*; PfENT1, *Plasmodium falciparum*; ScFUN26, *Saccharomyces cerevisiae*; AtENT1, *Arabidopsis thaliana*; NNT, *Naegleria* sp.; LmNnt1-1, *Leishmania major*; HsENT1/2, MmENT1, human and mouse ENT1/2; HsCNT1/2, human concentrative nucleoside transporters—used as the out-group). MUSCLE-aligned sequences were subjected to neighbor-joining analysis; node labels give bootstrap support from 1,000 replicates (values < 30% omitted). Scale bar, 0.5 substitutions per site. **(B)** Predicted tertiary structures: AlphaFold models of the four TgENT paralogues are shown in ribbon representation (top-view, upper row; side-view, lower row). Each adopts the canonical 11–13 trans-membrane helix bundle typical of ENT family members, with the central permeation pore clearly visible in the side view. Colors match the gene names in panel A. **(C)** Quantitative structural comparison to human ENT1 (PDB 6OB7). Pore dimensions were measured along the long and short axes (Å) for each TgENT model and for HsENT1, and Cα-root-mean-square deviations (RMSDs) were calculated after SuperPose alignment to 6OB7. Heat-map shading reflects magnitude (scale bar, right). Cartoon underneath illustrates the pore-width measurements on the HsENT1 template. **(D)** Stage-specific expression of TgENT transcripts. Heat-map of log2-transformed TPM values (+1) extracted from the ToxoDB developmental transcriptome. EES 1–5, enteroepithelial stages in cat epithelium; sporulating oocysts; tachyzoites; tissue cyst bradyzoites; unsporulated oocysts ^19,20^. TgAT1 is highest in enteroepithilial and early tachyzoite stages, TgENT1 is broadly expressed, TgENT2 peaks during sporulation, and TgENT3 shows moderate expression throughout.

**Table 1.** BLAST results for *T. gondii* sequences with high similarity with Human ENT1 and *Plasmodium falciparum* ENT1.

| Gene ID | Name | Localization by hyperLOPIT<sup>36</sup> | PFam ID | PFam Description | Phenotype Score<sup>24</sup> |
| --- | --- | --- | --- | --- | --- |
| TGME49_233130 | TgENT3 | PM integral, Golgi | PF01733 | Equilibrative nucleoside transporter | 0.04 |
| TGME49_244440 | TgAT1 | PM integral, Golgi | PF01733 | Equilibrative nucleoside transporter | 1.02 |
| TGME49_288540 | TgENT1 | None | PF01733 | Equilibrative nucleoside transporter | -3.68 |
| TGME49_500147 | TgENT2 | None | PF01733 | Equilibrative nucleoside transporter | 0.47 |

### Nucleoside analogs growth assay

We hypothesized that the deletion of genes encoding nucleoside transporter homologs would negatively affect parasite growth. However, none of the single knockout strains displayed significant differences in either short-term or long-term growth rates compared to the parental strain (**Fig. S2**). The growth phenotype of TgENT1 knockdown parasites is described separately below (**Fig. 4**).

Among the four ENT homologs in *Toxoplasma gondii*, only TgAT1 has been biochemically characterized^15^. To determine whether deletion of TgENT2, TgENT3, or TgAT1 alters parasite sensitivity to the toxic nucleoside analogs 9-β-D-arabinofuranosyladenine (Ara-A)^18^ and 5-fluorouracil (5-FU)^21^, we grew the respective knockout strains in 96-well plates for three days in the presence of increasing concentrations of each analog. Infected monolayers were then fixed and stained with a rabbit anti-*Toxoplasma* (anti-Tg) antibody. Using immunofluorescence microscopy, we measured the area of at least 50 parasitophorous vacuoles (PVs) per condition; the mean PV size served as a quantitative read-out of growth inhibition (**Fig. 2**).

**Figure 2.**
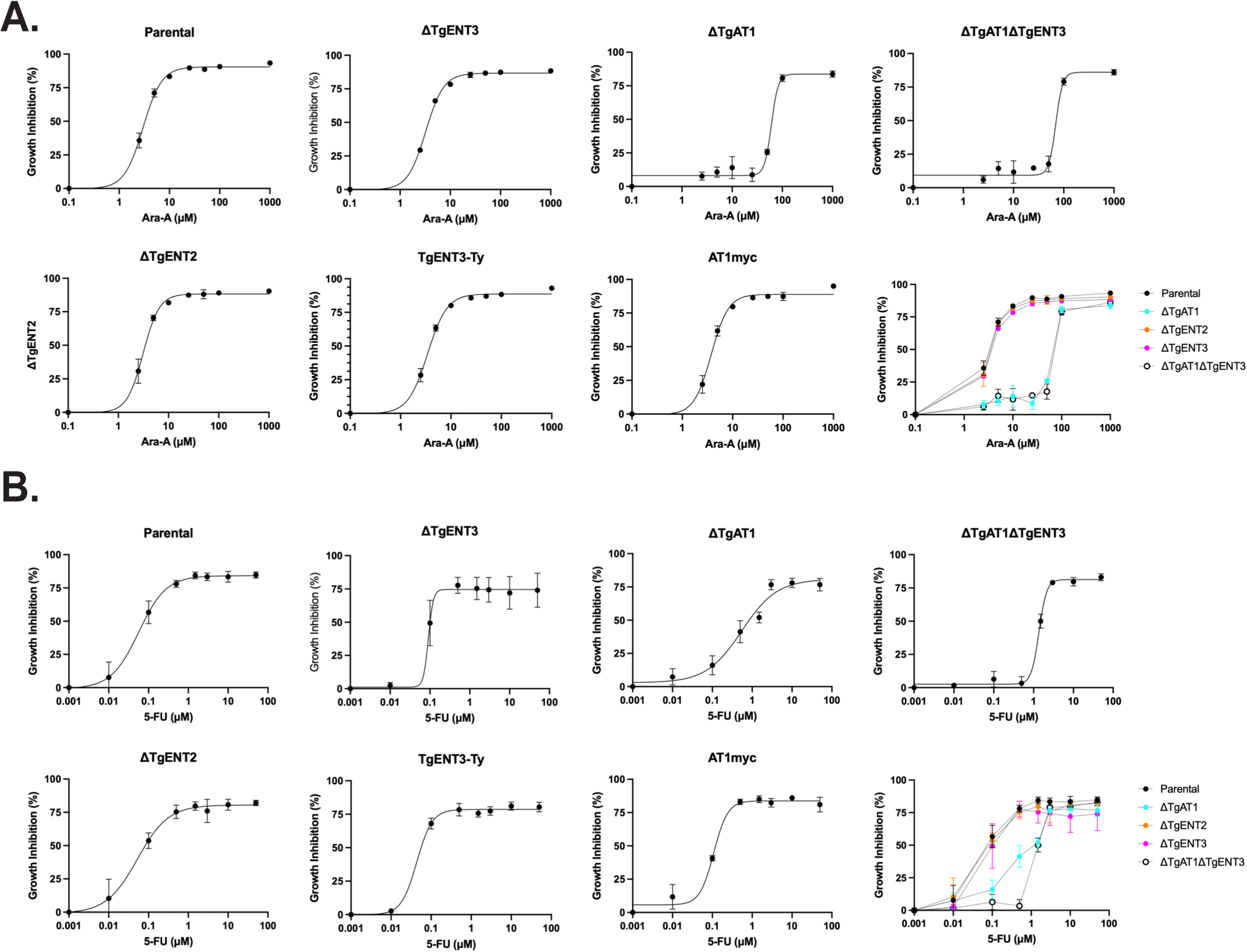
Deletion of TgAT1 and TgENT3 alters *Toxoplasma* sensitivity to toxic nucleoside analogs. Parental ME49 parasites and the indicated mutants were allowed to replicate for 72 h in HS27 monolayers in the presence of increasing concentrations of **(A)** the adenosine analog 9-β-D-arabinofuranosyl-adenine (Ara-A) or **(B)** the pyrimidine analog 5-fluorouracil (5-FU) Cultures were fixed, stained with anti-*Toxoplasma* (anti-Tg) antibody, and the mean area of ≥50 parasitophorous vacuoles (PVs) per condition was measured by automated image analysis. Growth inhibition (% ± SEM) was calculated relative to drug-free controls and fitted to a four-parameter logistic curve (GraphPad Prism). Data are from three biological replicates; each performed in technical triplicate. Panel layout: For each drug, individual dose–response curves are shown for the parental line, single knockouts (ΔTgENT3, ΔTgENT2, ΔTgAT1), the double knockout (ΔTgAT1ΔTgENT3), and the corresponding epitope-tagged complementation lines (ΔTgENT3::TgENT3-Ty1, ΔTgAT1::TgAT1-myc). The right-most graph in each row overlays all strains for direct comparison (color key, inset).

Consistent with TgAT1’s role as a purine transporter^15^, ΔTgAT1 parasites showed increased resistance to the adenosine analog Ara-A (**Fig. 2A**). Unexpectedly, ΔTgAT1 also exhibited resistance to the pyrimidine analog 5-FU (**Fig. 2B**), suggesting broader substrate specificity than previously recognized. In contrast, the ΔTgENT2 strain did not differ from the parental line (**Fig. 2**), likely because TgENT2 is not expressed during the tachyzoite stage (**Fig. 1D**). Similarly, the ΔTgENT3 single knockout showed no difference in sensitivity relative to the parental line under drug treatment (**Fig. 2**). However, the double knockout ΔTgAT1ΔTgENT3 formed significantly larger PVs than ΔTgAT1 under 10 μM Ara-A, indicating that deletion of TgENT3 further enhances resistance to this analog. Conversely, ΔTgAT1ΔTgENT3 vacuoles were significantly smaller than those of ΔTgAT1 under 5 μM 5-FU, although they remained larger than parental vacuoles. Collectively, these observations suggest that while TgENT3 deletion may confer increased resistance to Ara-A in the ΔTgAT1ΔTgENT3 background, it also leads to heightened sensitivity to 5-FU.

### Role of TgENTs homologs in chronic infection

We successfully generated knockout strains for TgAT1, TgENT2, and TgENT3 using CRISPR/Cas9-mediated gene disruption in the ME49 background. Multiple attempts to generate a TgENT1 knockout strain were unsuccessful, suggesting this gene may be essential for parasite viability. This prompted us to instead use the mini auxin-inducible degron (mAID) system for conditional knockdown analysis (**Fig. 4**) in the RH background.

Based on published transcriptomic data^19,20^ (**Fig. 1D**), TgENT1, TgENT3, and TgAT1 are constitutively expressed throughout the parasite life cycle, whereas TgENT2 expression is restricted to bradyzoite and sexual developmental stages, explaining the lack of phenotype in tachyzoite growth assays. Given this distinct expression pattern, as well as the unexpected lack of growth defects in single knockout strains, we next investigated potential changes in cyst formation using an *in vitro* and *in vivo* model.

We assessed the impact of TgENT deletion on parasite differentiation by culturing human foreskin fibroblasts (HS27) infected monolayers cultured in RPMI medium adjusted to pH 8.2 and atmospheric CO_2_. Monolayers were fixed after two, three, or seven days, and differentiation rates were quantified using immunofluorescence assays (IFA) with the lectin Dolichos Biflorus Agglutinin (DBA), which stains *T. gondii’s* cyst wall^22^. Parasites were co-stained with anti-Tg antibody, which identifies both tachyzoites and bradyzoites, to quantify all vacuoles in the monolayers and calculate differentiation ratios among strains. ΔTgAT1 and ΔTgAT1ΔTgENT3 parasites exhibited significantly slower differentiation. Both ΔTgAT1 and ΔTgAT1ΔTgENT3 have delayed differentiation, exemplified by lower number of DBA vacuoles in the population (**Fig. 3A**). Moreover, after 3 days in pH 8.2 medium, the DBA-positive vacuoles formed by ΔTgAT1 and ΔTgAT1ΔTgENT3 parasites are smaller than parental and complement strains (**Fig. S3**). By day 7, only 88% of ΔTgAT1 and 63% of ΔTgAT1ΔTgENT3 parasites were DBA-positive, compared to approximately 99% in the parental and ΔTgENT3 strains (**Fig. 3A**). As we noticed at day 3, the ΔTgAT1 and ΔTgAT1ΔTgENT3 DBA positive vacuoles are significantly smaller at day 7 as well compared to other strains (**Fig. 3B**). While ΔTgENT3 did not show a reduction in differentiation rate, the accentuated phenotype in the double deleted strain suggests an interplay between TgAT1 and TgENT3 in parasite differentiation under alkaline stress.

**Figure 3.**
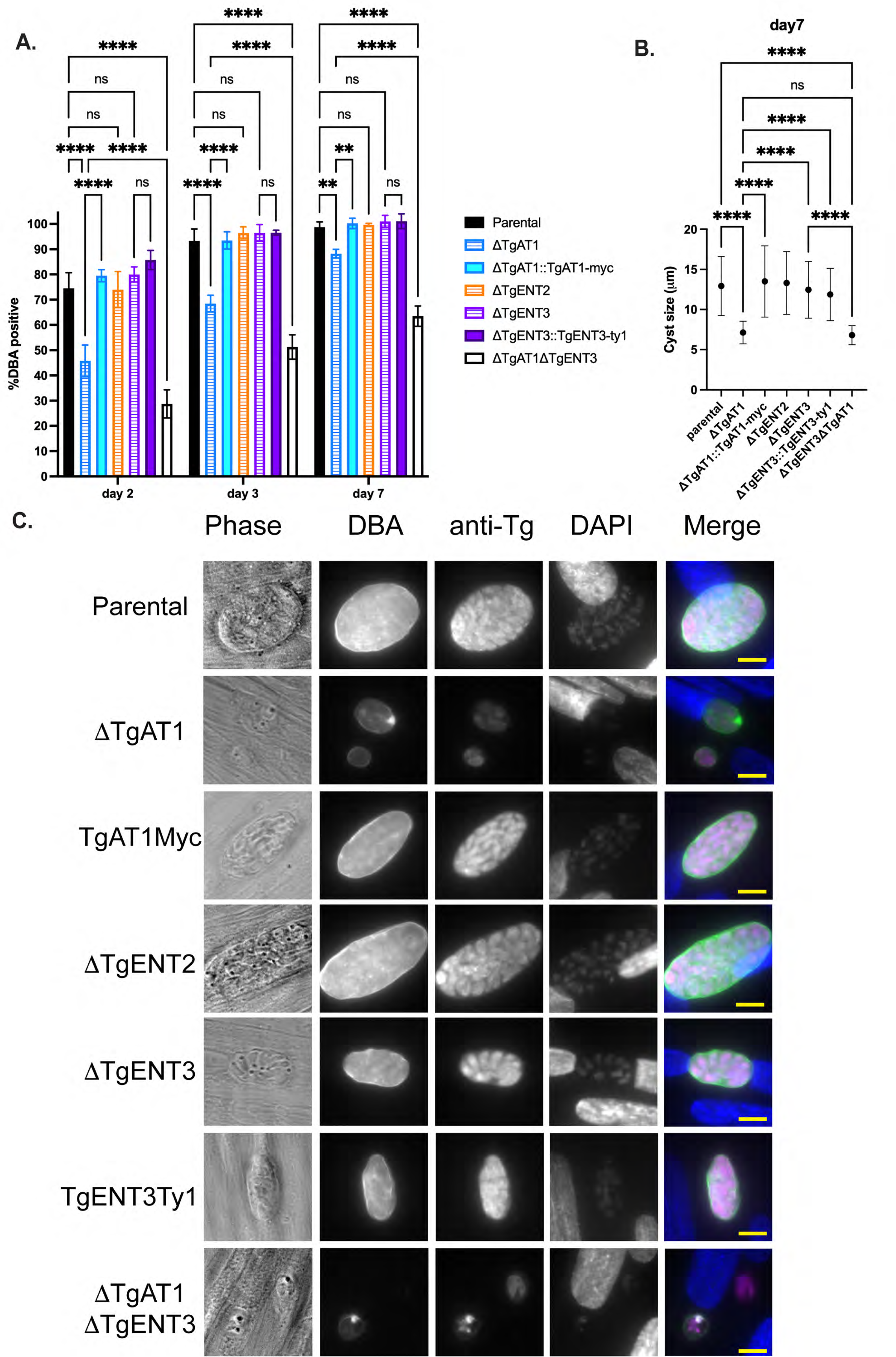
Simultaneous deletion of TgENT3 and TgAT1 delays differentiation. **(A)** Percentage of differentiation of the parental line, single knockouts (ΔTgENT3, ΔTgENT2, ΔTgAT1), the double knockout (ΔTgAT1ΔTgENT3), and the corresponding epitope-tagged complementation lines TgENT3-Ty1 (ΔTgENT3::TgENT3-Ty1) and TgAT1-myc (ΔTgAT1::TgAT1-myc) at day 2, 3, and 7 post invasion of HS27 monolayers cultured in alkaline stress conditions (pH 8.1 and atmospheric CO_2_). Parasites were stained with immunofluorescent antibodies DAPI, anti-Tg, and DBA. Bars show the percentage of DBA positive vacuoles (thresholds set based on parental line intensity) over total vacuoles (anti-Tg). Slides were imaged with BioTEK Cytation 7 and quantified using the Gen5 software (≥100 vacuoles were counted per replicate, n = 3 biological replicates). Statistical significance was assessed using one-way ANOVA, (****P < 0.0001, ns = non-significant). **(B)** Average sizes of each cyst (DBA positive vacuole) on day 7 post invasion. **(C)** Representative images of vacuoles from each parasite line. Scale bar, 10 µm

To assess long-term differentiation *in vivo*, male C57BL/6 mice were intraperitoneally infected with 250 parasites. Four weeks post-infection, mice were euthanized in accordance with Institutional Animal Care and Use Committee (IACUC) guidelines. Brains were extracted surgically, homogenized in PBS through sequential passage using 18-, 20-, and 22-gauge needles, fixed, and stained with DBA. Cyst numbers and sizes were quantified for each strain, as previously done^23^, (**Fig. S4**). This experiment was independently replicated twice, with four mice per strain in each replicate. No significant differences in cyst counts were observed among the strains (**Fig. S4A**). These findings suggest that, despite initially impaired differentiation observed in ΔTgAT1 and ΔTgAT1ΔTgENT3 parasites, these mutants ultimately compensate for this deficit during chronic infection in mice. The cyst sizes for ΔTgENT3 and ΔTgAT1ΔTgENT3 were significantly lower compared to all other strains, suggesting that TgENT3 may have important roles during bradyzoite replication, as cyst sizes are proportional to parasite growth (**Fig. S4B**).

To investigate whether the comparable tachyzoite growth rate (**Fig. S2**) observed in ΔTgAT1ΔTgENT3 was due to compensatory expression of another TgENT family member, we performed quantitative PCR (qPCR) analyses. We compared the expression levels of the remaining TgENT transporters, TgENT1 and TgENT2, across tachyzoites derived from ΔTgAT1ΔTgENT3, single knockout strains, and the parental strain. qPCR analysis revealed a ∼3-fold upregulation of TgENT1 transcripts specifically in ΔTgAT1ΔTgENT3 parasites compared to parental controls (p < 0.05), while TgENT2 expression remained unchanged (**Fig. 4A**). Notably, single knockout strains, ΔTgAT1 and ΔTgENT3, showed no compensatory changes in TgENT1 or TgENT2 (**Fig. 4A**). These data indicate that the absence of both TgAT1 and TgENT3 may trigger compensatory upregulation of TgENT1, maintaining tachyzoite growth rates similar to single knockout strains.

**Figure 4.**
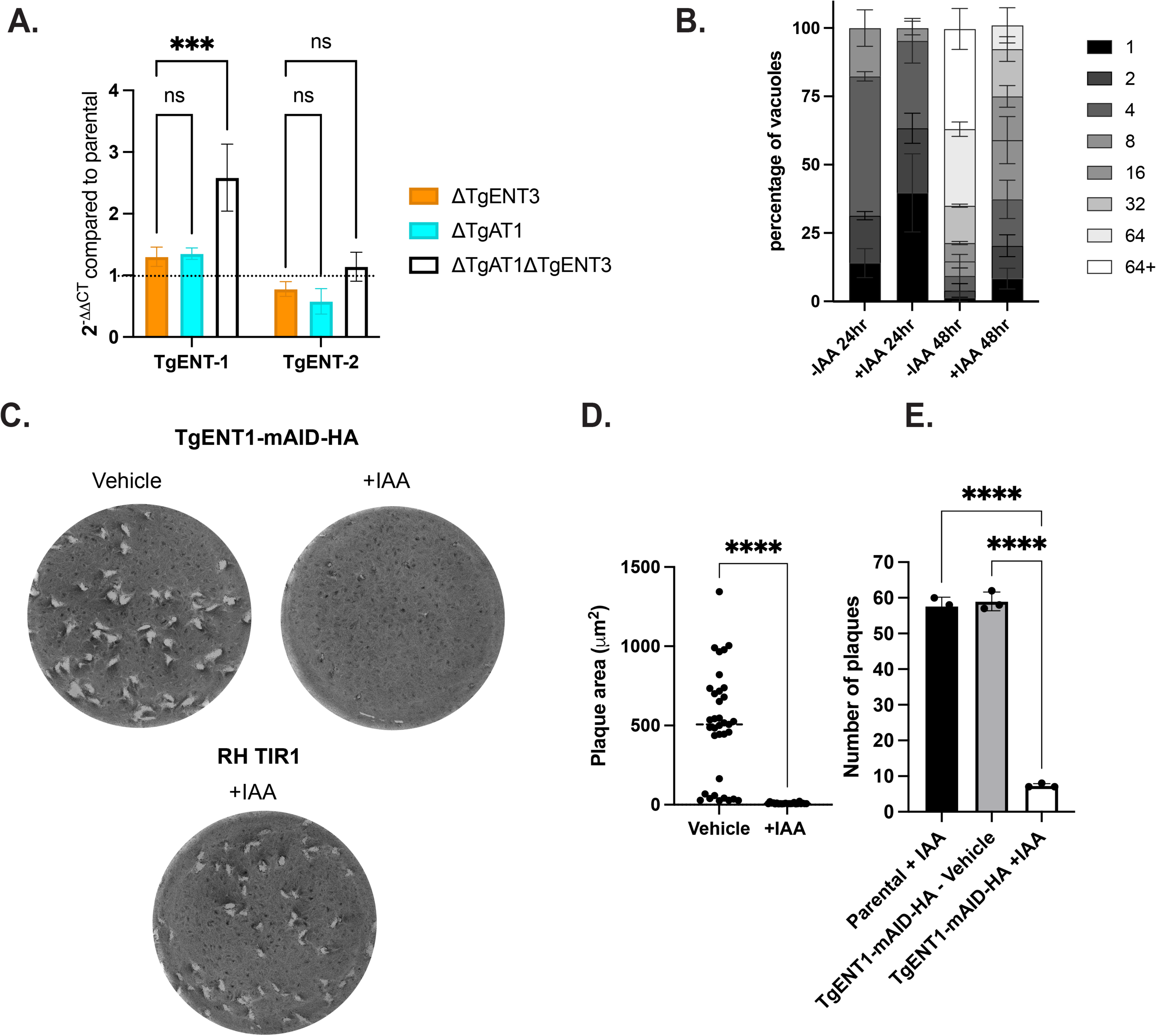
TgENT1 is transcriptionally upregulated in ΔTgAT1ΔTgENT3 parasites and is indispensable for intracellular replication and plaque formation. **(A)** qRT-PCR analysis of compensatory TgENT expression. Total RNA from tachyzoites of the indicated mutant lines was analyzed by qRT-PCR with primers specific for **TgENT1** and **TgENT2**. Bars show 2^ΔΔCt^ values (mean ± SEM, n = 3 biological replicates) normalized to the parental line (dotted line = 1). Only the double knockout (ΔTgAT1ΔTgENT3, open bars) exhibits a ∼3-fold induction of TgENT1 (one-way ANOVA with ***P < 0.001); TgENT2 is unchanged (ns). **(B)** Acute intracellular growth after TgENT1 depletion. TgENT1-mAID-HA parasites were pre-treated for 48 h with vehicle or 500 µM indole-3-acetic acid (+IAA) to trigger auxin-inducible degradation, then allowed to invade fresh HS27 monolayers for a further 48 h under the same conditions. Stacked bars depict the distribution of vacuoles containing 1, 2, 4, 8, 16, 32, 64 or >64 parasites (≥50 vacuoles counted per replicate; 3 replicates). **(C)** Plaque-formation assay. Wells were inoculated with 50 tachyzoites of the indicated strains and cultured for 9 days ± 500 µM IAA before fixation and crystal-violet staining. Representative wells are shown. **(D)** Quantification of plaque area. Individual plaque areas from TgENT1-mAID-HA wells (vehicle vs +IAA) are plotted (median ± inter-quartile range; n = 15–20 plaques). Student t-test, ****P < 0.0001. **(E)** Quantification of plaque number. Mean ± SEM plaque counts per well from three independent experiments. T-test, ****, P < 0.0001.

### TgENT1 conditional deletion results in complete growth arrest

TgENT1 has a markedly negative genome-wide CRISPR phenotype score of –3.68^24^. On this scale, increasingly negative values typically denote progressively stronger fitness costs, underscoring TgENT1’s critical importance for parasite survival (**Table 1**). To investigate the role of TgENT1, we developed a parasite line, TgENT1-mAID-HA, in RH backgrond with regulated TgENT1 expression via 3-indoleacetic acid (IAA) treatment ^25^. We assessed the impact of TgENT1 depletion on parasite growth by culturing the parasites with IAA for 48 hours. The treated parasites exhibited reduced replication when invading new HS27 monolayers (**Fig. 4B and C**), indicating that TgENT1 expression is crucial for normal growth and replication. Additionally, a long-term growth defect was observed over nine days of continuous culture, as evidenced by the formation of smaller and fewer plaques (**Fig. 4D**).

### TgENT1 partially colocalizes to the PLVAC

Using the TgENT1-mAID-HA construct, we were able to visualize the association of TgENT1 within *T. gondii* parasites. When cultured with 500 µM indole-3-acetic acid (IAA), we observed the disappearance of the TgENT1 signal, confirming the system functionality (**Fig. 5**). Surprisingly, TgENT1 localization differed markedly from typical plasma membrane-associated ENTs. Instead, HA-TgENT1 displayed a distinctive punctate pattern within the parasite cytoplasm, suggesting it is localized within intracellular organelles (**Fig. 5**).

**Figure 5.**
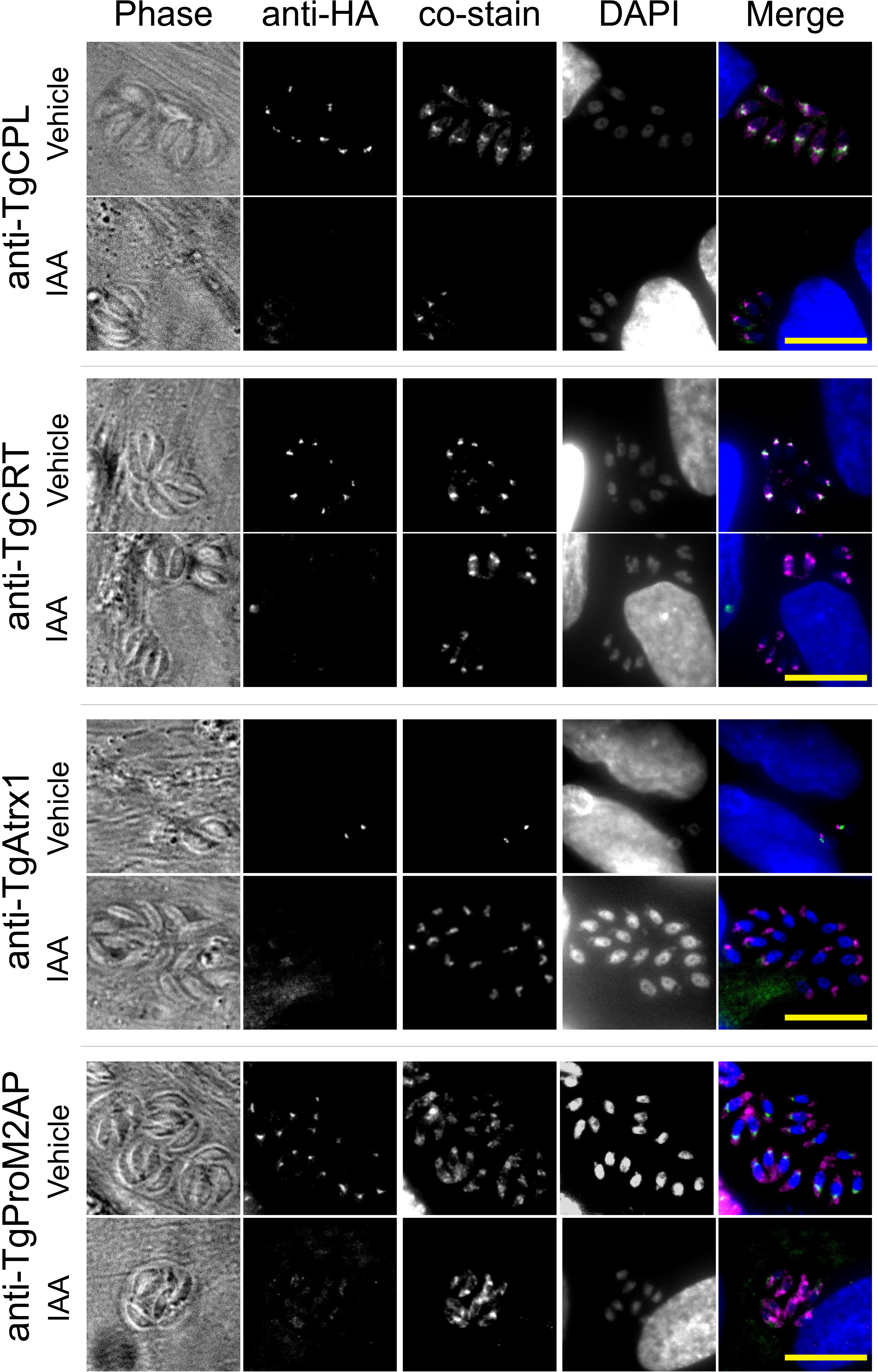
TgENT1 resides in the PLVAC and is eliminated by auxin-induced degradation. Representative immunofluorescence images of TgENT1-mAID-HA tachyzoites pre-treated for 48 h with vehicle or 500 µM indole-3-acetic acid (+IAA), then grown in fresh HS27 monolayers for a further 48 h under the same conditions. Parasites were fixed and stained with anti-HA (second column) plus one of the indicated compartment markers (third column): *T. gondii* cathepsin protease L (TgCPL) or *T. gondii* chloroquine-resistance transporter (TgCRT) for the PLVAC, TgAtrx1 for the apicoplast, and TgProM2AP for the endosome-like/microneme compartment. Columns show (left to right) phase-contrast, anti-HA, co-stain, DAPI, and the merged overlay (anti-HA = green, co-stain = magenta, DAPI = blue). Under vehicle conditions TgENT1-HA displays a punctate pattern that co-localizes with PLVAC markers TgCPL and TgCRT, but only weakly with TgProM2AP and not with TgAtrx1, consistent with predominant residence in PLVAC. Upon IAA treatment the HA signal is virtually abolished, confirming efficient auxin-inducible degradation. Scale bar, 10 µm.

To pinpoint the intracellular compartment that hosts TgENT1, we performed immunofluorescence assays (IFA) on parasites expressing HA-tagged TgENT1 alongside well-validated organelle markers. Co-staining was carried out with antibodies against cathepsin protease L (TgCPL) and the chloroquine-resistance transporter (TgCRT), both definitive markers of the plant-like vacuole (PLVAC)^26^, as well as the propeptide of microneme protein 2 (TgProM2AP), which labels the endosome-like compartment ^27^, and an anti-apicoplast (TgAtrx1) antibody^28^. Here, M(X) represents Manders’ colocalization coefficients which quantify the fraction of antibody X’s fluorescence that shares the same pixels as the HA signal (1.0 = complete overlap; 0 = none). Across all markers, in control conditions, the TgENT1 HA-tagged protein shows partial colocalization with PLVAC markers: TgCPL (Pearson’s r =0.84; M(CPL) =0.90) and TgCRT (Pearson’s r =0.79; M(CRT) =0.54), weak overlap by the microneme protein TgProM2AP (Pearson’s r =0.61; M(M2AP) =0.09), and virtually no overlap with TgAtrx1(Pearson’s r =0.10; M(Atrx1) =0.05). After IAA-induced depletion, Manders’ coefficients collapse to near background (TgCPL 0.08; TgCRT 0.02; TgProM2AP 0.09; TgAtrx1 0.00) and Pearson’s r values likewise fall (CPL 0.21; CRT 0.03; ProM2AP 0.25; Atrx1 0.05), indicating loss of specific HA signal and that the residual fluorescence no longer maps to these organelles. Manders and Pearson’s colocalization coefficients are based on three different biological experiments. Together, these results identify the PLVAC as the principal compartment for TgENT1.

### PLVAC measurement

Given the partial colocalization of TgENT1 and TgCRT and TgCPL (**Fig. 5**), we evaluated PLVAC morphology following TgENT1 depletion by performing knockdown experiments in replicating parasites 36 hours post-infection. Parasites were mechanically liberated and subsequently inoculated onto fresh HS27 monolayers for 30 minutes before fixation and staining. Importantly, TgENT1 knockdown parasites retained their ability to invade host cells efficiently. Immunofluorescence assays were employed to measure PLVAC dimensions using an anti-TgCPL antibody (**Fig. 6**). Area from 50 TgCPL puncta per experimental condition were measured in triplicate, and comparative statistical analyses were performed using t-test. Consistent with our hypothesis, TgENT1 depletion resulted in measurable swelling of the PLVAC, further implicating TgENT1 as important for maintaining PLVAC homeostasis.

**Figure 6.**
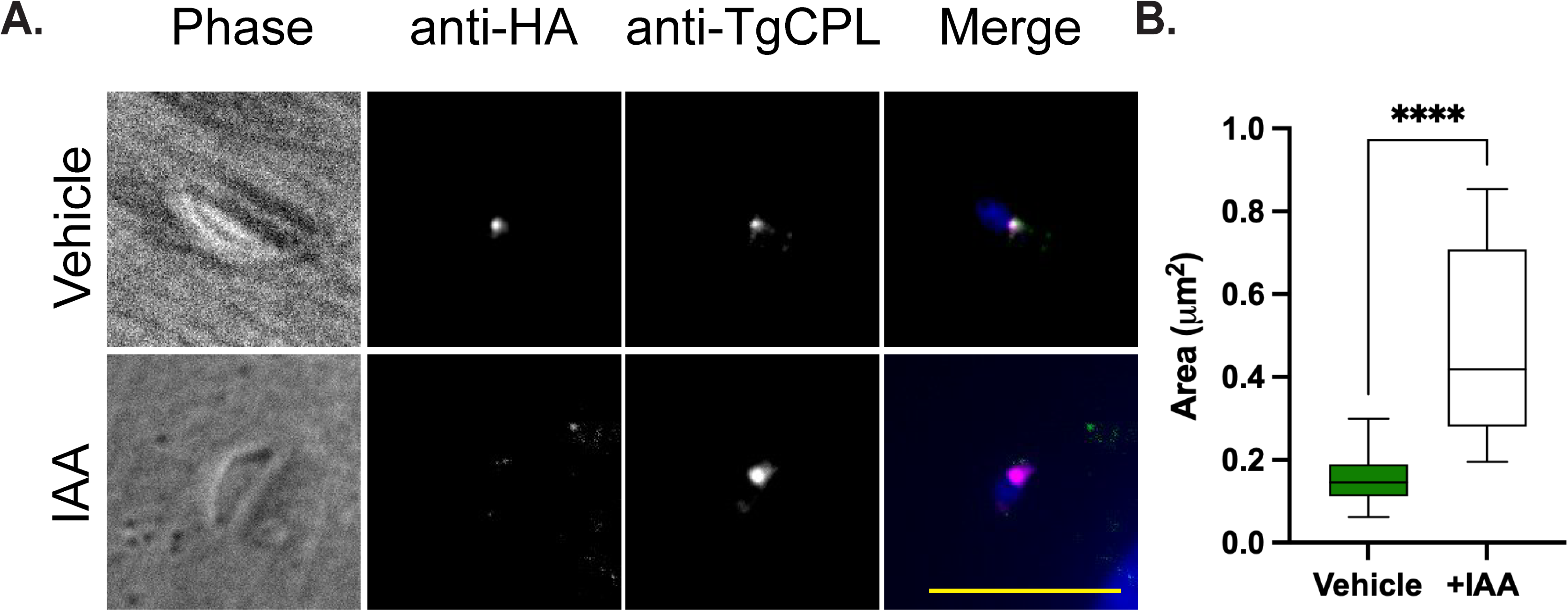
Loss of TgENT1 causes swelling of the PLVAC. **(A).** TgENT1-mAID-HA parasites were grown for 48 h post-infection, mechanically released, and allowed to reinvade new HS27monolayers for 30 minutes in the continued presence of vehicle or 500 µM IAA. Cells were fixed and co-stained with anti-HA (TgENT1, green) and anti-TgCPL (PLVAC marker, magenta); nuclei were counter-stained with DAPI (blue). TgENT1 is readily detected in the central PLVAC punctum under vehicle conditions but is efficiently depleted after auxin treatment, coincident with an enlarged TgCPL-positive compartment. Scale bar, 10 µm. **(B) Quantification of PLVAC size.** Area of 50 individual CPL-positive structures per condition (three independent experiments) was measured in FIJI. Box-and-whisker plots show median, inter-quartile range, and 5th–95th percentiles. Auxin-induced TgENT1 depletion increased PLVAC area nearly five-fold, student t-test, **** P < 0.0001, suggesting that TgENT1 activity is required to maintain normal vacuolar homeostasis.

### Transcriptomic analysis of TgENT1 knockdown

Transcriptomic profiling following conditional knockdown of TgENT1 revealed significant alterations in genes associated with purine metabolism, reinforcing TgENT1’s proposed function as a nucleoside transporter (**Fig. 7**). Principal-component analysis (**Fig. 7A**) cleanly separates IAA-treated and control tachyzoites along PC1 and shows a further time-dependent shift on PC2, underscoring a rapid and progressive transcriptional response to TgENT1 loss. An accompanying heat-map of (**Fig. 7B**) highlights the 50 most variable transcripts. Volcano plots of differentially expressed genes (**Fig. 7C**) display a broader and more pronounced response by 48 h, with the number and magnitude of regulated transcript expanding and purine-stress markers (e.g., nucleoside phosphatases and phosphodiesterases) among the most strongly up-regulated. For example, TGGT1_225290 (GDA1/CD39 nucleoside phosphatase) demonstrated progressive upregulation from day 1 (log₂ fold change = 2.98) to day 2 (log₂ fold change = 6.21). Similarly, TGGT1_259960 (putative nucleoside-diphosphatase) exhibited strong induction from day 1 (log₂ fold change = 2.81) to day 2 (log₂ fold change = 5.92), suggesting a sustained compensatory effort to restore purine homeostasis when nucleoside uptake is impaired. Importantly, other TgENT family members showed no significant expression changes.

**Figure 7.**
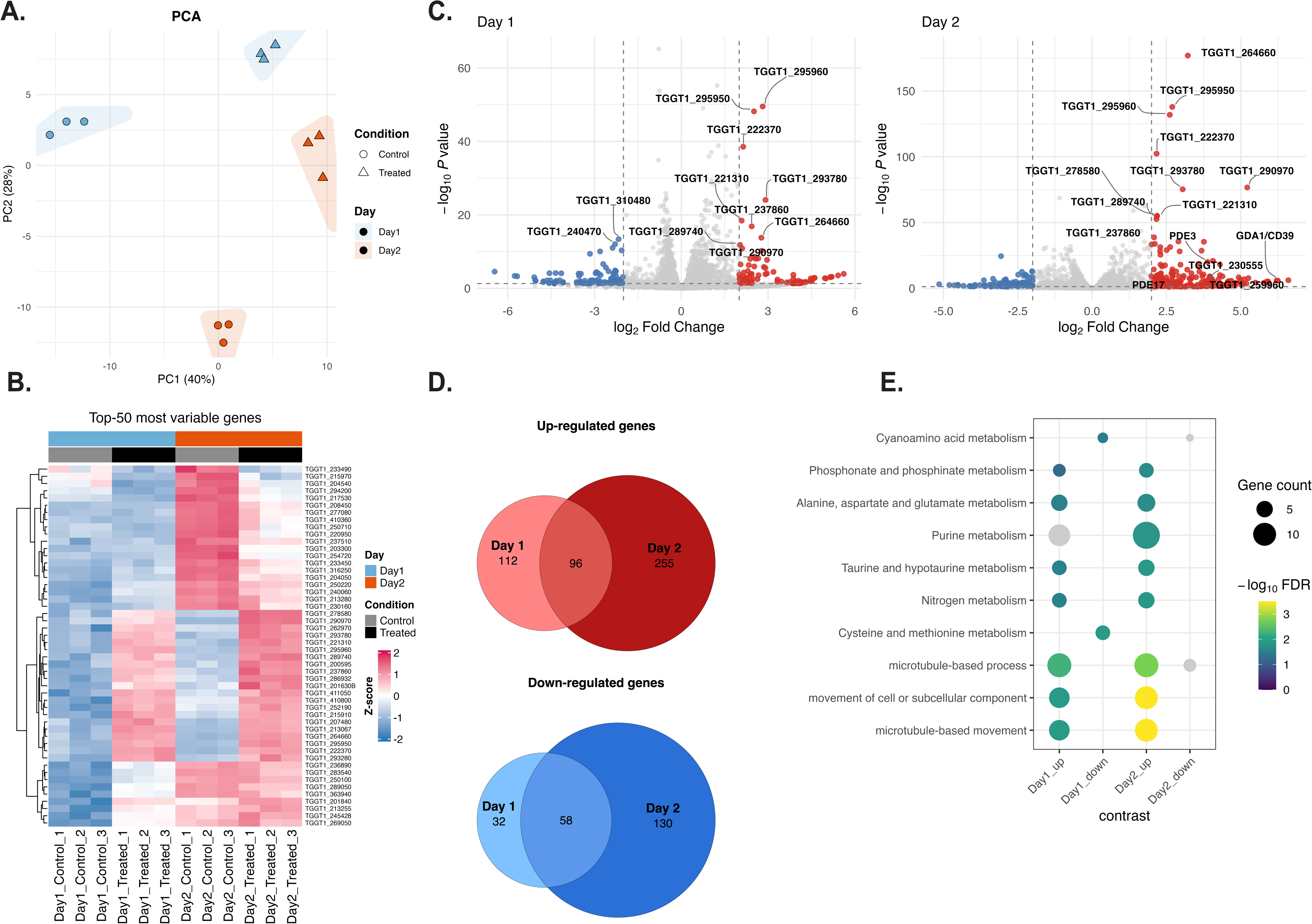
Transcriptomic response to acute TgENT1 depletion reveals a progressive, purine-centric stress program. TgENT1-mAID-HA tachyzoites (three biological replicates) were pre-treated for 48 h with vehicle or 500 µM IAA, then allowed to invade fresh HS27 monolayers under the same conditions and harvested 24 h (Day 1) or 48 h (Day 2) after invasion. Poly-(A) RNA was sequenced (Illumina NovaSeq; 150 bp PE), reads were mapped to the GT1 genome, and differential expression was analyzed with DESeq2 (|log₂FC| ≥ 1; FDR < 0.01). **(A) Principal-component analysis (PCA).** PC1 (40%) cleanly separates treated from control samples, whereas PC2 (28%) differentiates time points, indicating a robust, time-dependent transcriptional shift upon TgENT1 knockdown. **(B) Heat-map of the 50 most variable genes.** Z-scored expression values highlight a concerted up-regulation of purine- and nitrogen-metabolism genes (reds) and down-regulation of cytoskeletal / motility genes (blues) that intensifies from Day 1 to Day 2. **(C) Volcano plots of differentially expressed genes.** Pink, up-regulated; blue, down-regulated; grey, non-significant. The magnitude and number of responsive genes expand markedly by Day 2. The complete list of genes and fold change is presented of **supplemental table 4**. **(D) Overlap of DE genes between time points.** Venn diagrams show 96 genes up-regulated at both Day 1 and Day 2, and 58 genes consistently down-regulated, underscoring a core TgENT1-dependent regulon that is sustained and amplified over time. **(E) Functional enrichment analysis.** Dot-plot of the top Gene Ontology / KEGG terms (size = gene count; color = –log₁₀FDR). Up-regulated sets are dominated by **purine metabolism**, **nitrogen-compound metabolism**, and **cyano-/taurine-derivative pathways**, whereas down-regulated sets are enriched for **microtubule-based movement** and related processes, consistent with a shift from proliferation to metabolic stress adaptation.

In total, 208 transcripts were already altered at 24 h and this number more than doubled by 48 h, with 96 up- and 58 down-regulated genes shared between both time points (**Fig. 7D**), defining a core TgENT1-dependent regulon. Among the induced genes were several 3’5’-cyclic nucleotide phosphodiesterases (PDEs), notably PDE3 (TGGT1_233065; log₂ fold change = 1.30 at day 1, significantly increased to 3.88 at day 2), and PDE17 (TGGT1_257945; log₂ fold change = 2.92), suggesting heightened cyclic nucleotide turnover and signaling adaptation ^29^.

KEGG pathway analyses (**Fig. 6E**) revealed significant differential expression in pathways related to purine metabolism, nitrogen compound metabolism, and oxidoreductase activities. The enrichment of the purine metabolism pathway further intensified at day 2, according to KEGG pathway analysis (**Fig. 7E**) (fold enrichment = 2.39, p-value = 0.0008), featuring upregulated genes like TGGT1_230555^30^ (adenylate/guanylate cyclase; log₂ fold change = 3.84) and further emphasizing signaling changes upon TgENT1 knockdown. Additionally, TGGT1_290970^31^ (8-amino-7-oxononanoate synthase), involved in biotin biosynthesis, was substantially induced from day 1 (log₂ fold change = 2.57) to day 2 (log₂ fold change = 5.22), suggesting enhanced biotin-dependent metabolic processes.

Collectively, these transcriptomic findings suggest TgENT1’s central role in purine transport and homeostasis within *T. gondii*. The extensive and progressive compensatory responses triggered by TgENT1 knockdown, including purine salvage, metabolic reprogramming, and signaling adaptations, provide new insights into the parasite’s ability to adapt to disrupted nucleoside transport.

## Discussion

In this study, we have expanded the model of purine acquisition in *Toxoplasma gondii*, moving beyond a simple linear import pathway to reveal a dynamic, regulated network. HyperLOPIT spatial-proteomics has previously assigned both TgAT1 and TgENT3 to the parasite plasma membrane^36^ **(Table 1)**. Although our experiments did not verify this localization directly, the functional data reveal a broader principle of *metabolic resilience*: *Toxoplasma gondii* can rapidly re-wire its purine-uptake network to compensate for the loss of TgAT1 and TgENT3. Such plasticity is indispensable for an organism that is purine-auxotrophic and therefore wholly dependent on the host for nucleoside acquisition ^32^. Importantly, we also demonstrate that this adaptive capacity is limited during differentiation (**Fig. 3**). Understanding the mechanics, function, and ultimate failure point of this network provides unprecedented insight into the metabolic strategies governing the parasite’s lifecycle.

Our data show that deletion of the TgAT1 and TgENT3 triggers specific transcriptional upregulation of TgENT1 (**Fig. 4A**), a transporter that our localization studies showed to partially localize in the Plant-Like Vacuolar Compartment (PLVAC) (**Fig. 5**). These findings support a model where the parasite mobilizes an alternative salvage pathway. This pathway might rely on acquiring purines from the degradation of host-derived nucleic acids within the acidic, digestive environment of the PLVAC^26^ or from the recycling of its own nucleic acids, a function consistent with the PLVAC’s role in trafficking and catabolism ^26^. The adaptive function of the PLVAC and TgENT1 creates a crucial lifeline that sustains a baseline of purine homeostasis.

While this adaptive axis provides metabolic flexibility for tachyzoites, its capacity is finite, and its functional limit is most profoundly revealed during the critical developmental transition to the chronic stage. The failure of the ΔTgAT1ΔTgENT3 strain to develop as bradyzoites *in vitro* (**Fig. 3A and C**, and **S3**) likely stems from two interconnected deficiencies: a substrate and a signaling deficit. The substrate deficit may deplete critical precursors for cyst formation, including the sugar-nucleotide donors uridine diphosphate N-acetyl-D-glucosamine (UDP-GlcNAc) and uridine diphosphate N-acetyl-D-galactosamine (UDP-GalNAc). These activated sugars supply N-acetylglucosamine and N-acetylgalactosamine residues that decorate cyst-wall glycoconjugates and are important for proper wall assembly ^33^. More critically, however, this purine starvation likely precipitates a failure in cellular signaling. Purine salvage is the sole source of the adenosine required for ATP synthesis. ATP, in turn, is the exclusive substrate for adenylyl cyclases that generate cyclic AMP (cAMP), a master second messenger in *T. gondii*. The cAMP signaling pathway is known to be an important regulator of stage conversion, with fluctuations in cAMP levels acting as a key signal for maintaining the tachyzoite state or initiating differentiation into bradyzoites ^34^. Therefore, we propose that the ΔTgAT1ΔTgENT3 strain, while receiving the external cues to differentiate (e.g., alkaline stress), is metabolically incapable of mounting the appropriate cAMP signaling response required to execute this complex developmental program. Interestingly, although ΔTgAT1ΔTgENT3 mutants form smaller cysts than parental strains in mice, the total cyst burden remains unchanged, suggesting that the 30-day *in vivo* period allows the parasites to overcome the differentiation delay observed *in vitro* (**Fig. S4**).

Our study illuminates a clear transcriptional and phenotypic signature of nucleoside starvation in the absence of TgENT1, and several open questions make the next experimental steps especially compelling. First, the robust up-regulation of TgENT1 is currently inferred to reflect enhanced host-nucleoside scavenging; confirming this hypothesis with direct radiolabeled-substrate uptake assays will test our model and establish quantitative benchmarks for future transporter engineering. Second, the nucleoside-depletion phenotype reported for Δgra14 parasites, which are defective in parasite endocytosis of material from the cytosol of infected cells, suggests that the PLVAC has a central role in nucleoside salvage and invites a systematic dissection of each vacuolar effector in that pathway ^35^. Finally, untargeted metabolomics coupled to parallel measurements of high-energy substrates such as ATP and GTP and their signaling derivatives would provide a systems-level view of how disrupted uptake reverberates through the parasite’s metabolic and regulatory networks.

Ultimately, by deconstructing this purine acquisition network, we have revealed a dynamic interplay between metabolic adaptation and developmental signaling. The biological consequence of this network’s failure is unambiguous. By uncovering a scenario where the parasite’s adaptive capacity breaks down at the nexus of metabolism and developmental signaling, we have exposed a significant vulnerability. This metabolic bottleneck, where purine insufficiency hypothetically precipitates a signaling failure that blocks the establishment of chronic infection, represents a highly promising and conceptually novel target for therapeutic intervention. Further investigation offers the potential to develop strategies aimed specifically at disrupting the lifelong persistence that is the clinical hallmark of toxoplasmosis.

## Methods

### Parasite Maintenance

*Toxoplasma gondii* tachyzoites, ME49 and RH strains, were propagated in human foreskin fibroblasts (HS27; ATCC CRL-1634) at 37 °C in DMEM supplemented with 10 % FBS, 100 U/ml penicillin, and 100 µg/ml streptomycin. Cells and parasite lines were tested for mycoplasma monthly using a PCR test (Boca Scientific).

### Plasmid Construction

Final sequences are listed in **Supplementary Table 2**. PCRs employed Q5 High-Fidelity (long fragments) or Taq DNA polymerase (routine amplifications). Amplicons and vector backbones were assembled with Gibson Assembly (NEB) unless noted.

- **ΔTgENT3 donor** – synthetic plasmid (GenScript).
- **ΔTgENT3::TgENT3-Ty1 donor** – 23-DD plasmid digested with BglI. Synthetic insert (B1) ordered and inserted (Twist Bioscience).
- **ΔTgAT1 donor** – ΔTgENT3 donor digested with AvrII/NotI (insert) and SpeI/AflII (backbone); 5′ and 3′ TgAT1 homology arms amplified with P1/P2 and P3/P4.
- **TgENT3-DHFR-KO donor** – DHFR cassette (Addgene #80329) amplified with P5/P6 and inserted into ΔTgENT3 backbone (NotI/EcoRV).
- **mAID-HA-TgENT1 donor** – synthetic plasmid (GenScript).
- **ΔTgAT1::TgAT1-myc donor** – 24-DD plasmid digested with BssHI/ScaI. Synthetic inserts (B2 and B3) ordered and inserted (Twist Bioscience).
- **ΔTgENT2** – ΔTgENT3 donor digested with AflII/SnaBI. Synthetic insert (B4) ordered and inserted (Twist Bioscience).
- **Guide RNA plasmids** – pSS013 vector (Addgene) was BsaI-linearised; annealed IDT oligos were inserted by Gibson Assembly, transformed into *E. coli* DH5α, sequence-verified, and prepared with GeneJET Maxi kits.

### Genomic Editing and Transfection

Knock-outs, knock-ins, and mAID tagging were generated with dual-guide CRISPR–Cas9 as described (voltage 1500 V, 0.1 ms, two pulses; BTX ECM™ 830, 4-mm cuvette). Parasites harvested from confluent T25 flasks were washed once in cytomix (120 mM KCl, 0.15 mM CaCl₂, 10 mM K₂HPO₄/KH₂PO₄ pH 7.6, 25 mM HEPES, 2 mM EGTA, 5 mM MgCl₂). Electroporation mixtures contained 50 µg linearized donor DNA and 25 µg of each sgRNA plasmid in 400 µl cytomix. Parasites were immediately added to fresh HS27 monolayers. Drug selection began 24 h post-transfection (40 µM chloramphenicol or 3 µM pyrimethamine). Drug-resistant populations were single-cell-sorted (or cloned by limiting dilution), expanded, and validated by diagnostic PCR, Western blot, and immunofluorescence (primer sets P7– P28; see Supplementary Table 2).

### AlphaFold Modeling

Predicted protein structures were retrieved from the AlphaFold Protein Structure Database (AlphaFold DB) using their UniProt accession code. These predicted structures were aligned with human ENT1 (PDBID: 6OB7) using PyMOL version 3.1.4.1 (Schrodinger). Root Mean Square Deviation (RMSD) of each transporter relative to human ENT1 was calculated using PyMOL. Atomic measurements of pore width were also performed in PyMOL. As shown using HsENT1 in **Fig. 1C**, pore width was measured across the widest and narrowest points (yellow and red dashed lines). TgENT1, TgENT2, TgENT3, and TgAT1 pores were measured in the same manner.

### Immunofluorescence Assay

Cells were fixed in 3.7% formaldehyde for 20 minutes, washed twice with 1x PBS, blocked and permeabilized with 3% BSA and 0.2% Triton X-100 for 2 hours at room temperature (or, alternatively, overnight at 4°C). Primary antibodies were diluted 1:500 (unless otherwise stated in Supplemental Table 3) in blocking/permeabilization solution and incubated for 1 hour at room temperature (or overnight at 4°C). Cells were washed twice with 1x PBS, secondary antibodies were applied at 1:500 dilution (DAPI at 1:200) in blocking/permeabilization solution for one hour at room temperature, washed three times for 5 minutes, and glass coverslips were mounted and sealed.

### Pearson and Manders’ colocalization coefficients

Pearson’s *r* and Manders’ coefficients assess spatial overlap between two fluorescence signals after background removal. Pearson’s correlation coefficient (*r*) measures the linear proportionality of intensities on a pixel-by-pixel basis (−1 to 1), whereas Manders’ M1 and M2 report the fraction of fluorophore A (or B) signal that coincides with the other channel independent of intensity proportionality. Colocalization was quantified in raw 16-bit TIFF images using Fiji (ImageJ) Coloc2. After identical acquisition (no saturation; sequential capture to avoid bleed-through), images were opened without gamma adjustment, a uniform rolling-ball background subtraction was applied to both channels, and no nonlinear filtering or manual thresholding was performed. Reported values in the manuscript are the averages of 50 vacuoles per condition pooled from three independent biological experiments.

### Bradyzoite *in vitro* differentiation

The *in vitro* differentiation was performed as previously ^34^. Confluent HS27 fibroblast monolayers on glass coverslips were infected with freshly egressed tachyzoites (MOI ≈1; 2 h invasion + gentle wash; total pre-induction incubation 4 h). Cultures were then shifted to pre-warmed alkaline induction medium (RPMI 1640 supplemented with 1% heat-inactivated FBS, 50 mM HEPES, pH 8.20 ± 0.05 at 37 °C) and incubated in ambient air (≈0.04% CO₂) in a humidified chamber (no external CO₂) to maintain pH ≥8.0. Induction medium was replaced every 24 h with freshly adjusted medium; pH was spot-checked at each change. After 2-, 3-, or 7-days post-infection, coverslips were fixed (4% paraformaldehyde/PBS, 20 min), permeabilized (0.2% Triton X-100), and dual-stained with rabbit anti-*Toxoplasma* antibody (Thermo Fisher; 1:500) to delineate vacuoles and Dolichos biflorus agglutinin (DBA; 2–5 µg/ml) to label the cyst wall. A vacuole was scored “differentiated” if ≥75% of its perimeter showed a continuous DBA rim. For each condition ≥100 vacuoles per coverslip were scored blindly (≥3 biological replicates).

### Tachyzoite Intracellular Growth and Plaque Assays

Confluent HS27 (human foreskin fibroblast) monolayers on glass coverslips were infected with each *T. gondii* strain at an MOI yielding mostly isolated vacuoles (typically MOI 0.5–1.0) in standard pH 7.4 medium (DMEM + 10% FBS, 5% CO₂). After allowing invasion for 2 h at 37 °C, extracellular parasites were removed by two PBS washes and fresh medium was added. At the indicated post-infection time point in each experiment, cells were fixed in 4% paraformaldehyde (15 min), permeabilized (0.1% Triton X-100, 10 min), and blocked (3% BSA/PBS, 30 min). Parasites were labeled with anti-*Toxoplasma* (followed by Alexa Fluor–conjugated secondary antibody). For each biological replicate, 100 intact, non-overlapping parasitophorous vacuoles were randomly selected using a Cytation 7 imager (Agilent) under consistent exposure settings, and the number of tachyzoites per vacuole was recorded. Three biological replicates were performed per experiment. Vacuoles at the edge of fields, obviously disrupted, or with uncertain boundaries were excluded during counting.

For **plaque assays**, confluent HS27 monolayers in 6-well plates were infected with 1,000 of freshly egressed tachyzoites in 2 mL of complete medium. At endpoint, monolayers were fixed in 100% ethanol (10 min) and stained with 0.1% crystal violet (in 20% methanol) for 15–20 min, rinsed, and air-dried. Plaques (zones of host cell clearance) were imaged, counted manually and plaque areas were measured after background subtraction and uniform threshold application. At least three biological replicates (independent parasite preparations on different days) per strain were analyzed. Statistical analysis used one-way ANOVA with appropriate multiple comparison correction. **Quality controls.** Parasite viability and inoculum accuracy were confirmed by counting extracellular tachyzoites with a hemocytometer.

### RNA extraction, mRNA sequencing and transcriptomics analyses

Total RNA was extracted from *T. gondii* infected cells using the GeneJET RNA Purification Kit (Thermo Fisher Scientific) according to the manufacturer’s protocol. RNA quality and concentration were confirmed, and poly(A)-selected mRNA libraries were subsequently prepared by Novogene. Libraries were sequenced on an Illumina NovaSeq X Plus platform, generating 150-bp paired-end reads. All experiments were performed in biological triplicate, and the resulting data are publicly available in the NCBI Sequence Read Archive under BioProject accession PRJNA1247533. Reads were adapter- and quality-trimmed with Cutadapt, mapped to the reference genome using STAR in splice-aware mode, and uniquely aligned reads were summarized to gene-level counts with featureCounts. Differential expression was then evaluated in R with DESeq2, which applied median-of-ratios normalization, negative-binomial GLM fitting, and Benjamini– Hochberg correction to identify significantly regulated genes.

### Immunofluorescence Assay for PLVAC Morphology

PLVAC morphology was assessed 36 h post-infection in TgENT1 AID-HA tachyzoites ± IAA. Parasites were mechanically released (27-g needle) from host cells and immediately inoculated onto fresh confluent HS27 monolayers for a 30-min invasion pulse (37 °C). Cultures were fixed in 4% paraformaldehyde (20 min, RT) and stained with primary anti-TgCPL antibody (PLVAC marker) followed by appropriate fluorophore-conjugated secondary antibodies and DAPI. Imaging was performed under identical acquisition settings across conditions. For each condition, the cross-sectional area of the first 50 TgCPL-positive PLVAC puncta associated with intracellular parasites was quantified (Fiji) in each of three biological replicates (n = 50 vacuoles/condition). Areas were compared by t-test.

### Animal experiments

All animal work was approved by the University of Vermont IACUC (protocol PROTO202100038) and performed in accordance with institutional guidelines. Mice were housed under a 12 h light/12 h dark cycle with daily health monitoring that began one week before infection and continued until study end; animals were euthanized if predefined humane endpoints were met.

Two independent experiments were conducted. In each, four adult male CBA/J mice (The Jackson Laboratory) per parasite strain received an intraperitoneal inoculation of 250 freshly-egressed tachyzoites suspended in 200 µl sterile PBS. At 30 days post-infection (dpi), mice were euthanized following IACUC protocol, and whole cortices were promptly harvested for downstream analyses.

Brain immunofluorescence was conducted as previously reported ^37^. Briefly, the tissue was homogenized using serial needles passing through 18-, 20-, and 22-gauge needles, pelleted at 1000xg for 5 minutes, and fixed with 3.7% formaldehyde for 20 minutes. Pellets were washed twice with 1x PBS, blocked overnight in 1% BSA and 0.2% Triton X-100 in 1x PBS, primary antibodies diluted 1:500 were incubated overnight, washed twice with 1x PBS, secondary antibodies (1:500 dilution) and DAPI were added for 1 hour, washed twice with 1x PBS, and imaged using a microscope.

Cyst stained with DBA were wet mounted onto coverslips and scanned for DBA using a Cytation 7 imager (Agilent). 50 randomly found cysts per sample were measured and recorded using Agilent BioTek Gen5 software.

### qPCR

RNA was extracted from freshly lysed tachyzoites using the GeneJet RNA Purification Kit. Complementary DNA (cDNA) was subsequently synthesized employing the Maxima H Minus First Strand cDNA Synthesis Kit. Quantitative PCR (qPCR) reactions were then set up with primers Q1 – Q8 using the PowerSYBR Green PCR Master Mix.

### Statistical Analysis

GraphPad Prism 9 was used for all statistics. Data are mean ± SEM of ≥3 independent experiments. Group differences were analyzed by one-way ANOVA or unpaired two-tailed Student’s *t*-test (as indicated). *P* < 0.05 (\**), 0.01 (**), and 0.0001 (\*\*\*\**) were considered significant.

## Acknowledgments

This research was supported by a research grant from National Institutes of Health (P20 GM125498). We thank Dr. Harry de Koning for helpful discussions.

## Notes

### Competing Interest Statement

The authors have declared no competing interest.

### Summary of Updates

This revision includes data from TgENT2 mutant, which reviewers requested. Also, it has transcriptomics data for TgENT1.

