## Supplementary figures and images for "Impact of Equilibrative Nucleoside Transporters on *Toxoplasma gondii* Infection and Differentiation"

### Supplemental Figure 1

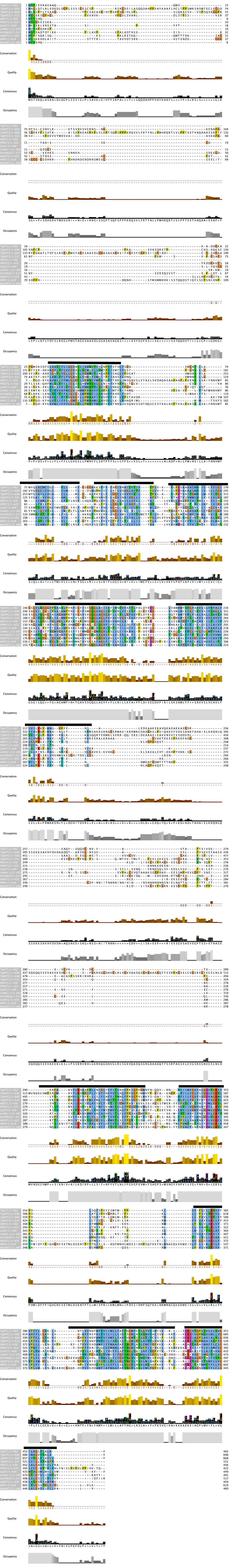

### Supplemental Figure 2

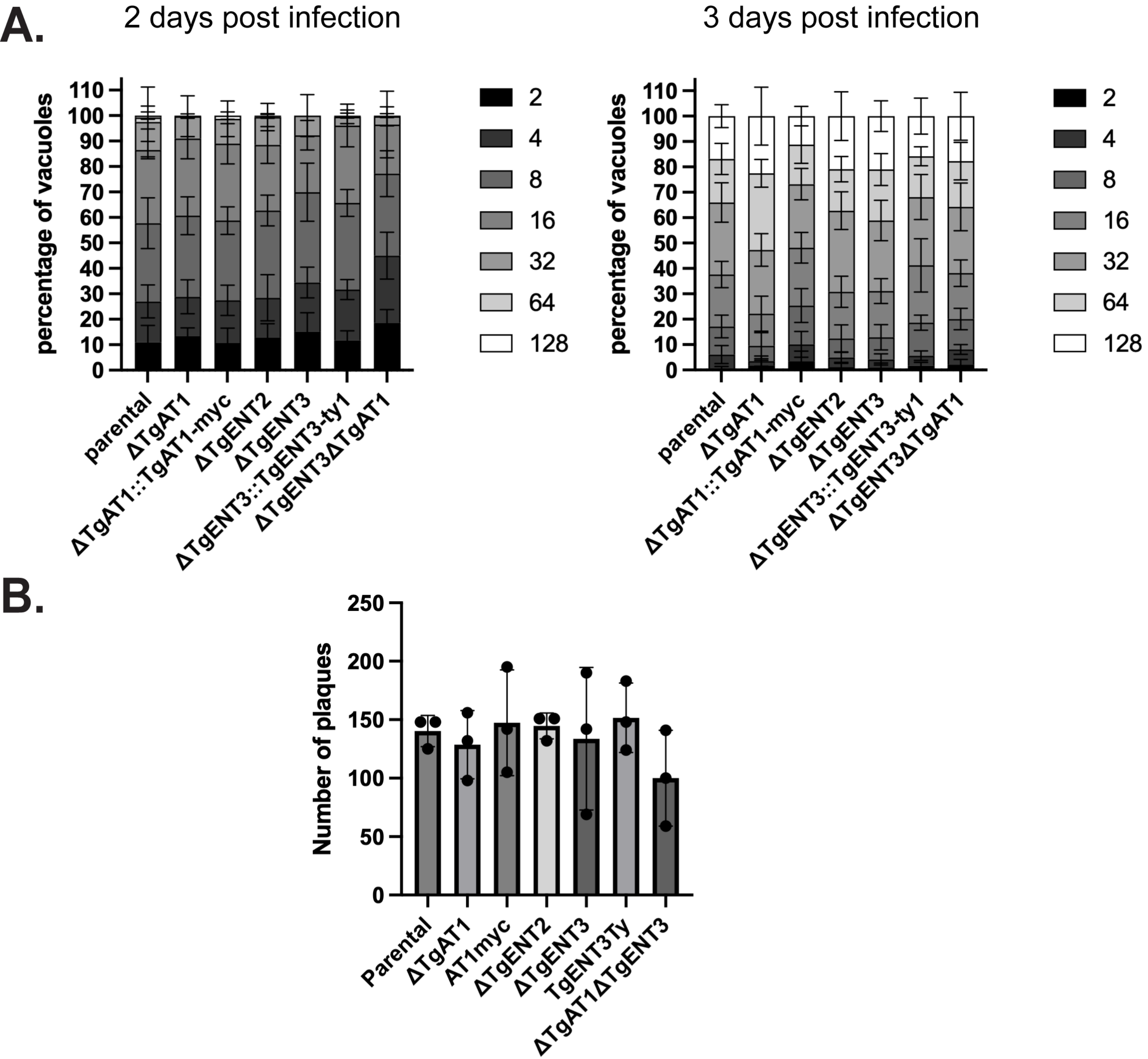

### Supplemental Figure 3

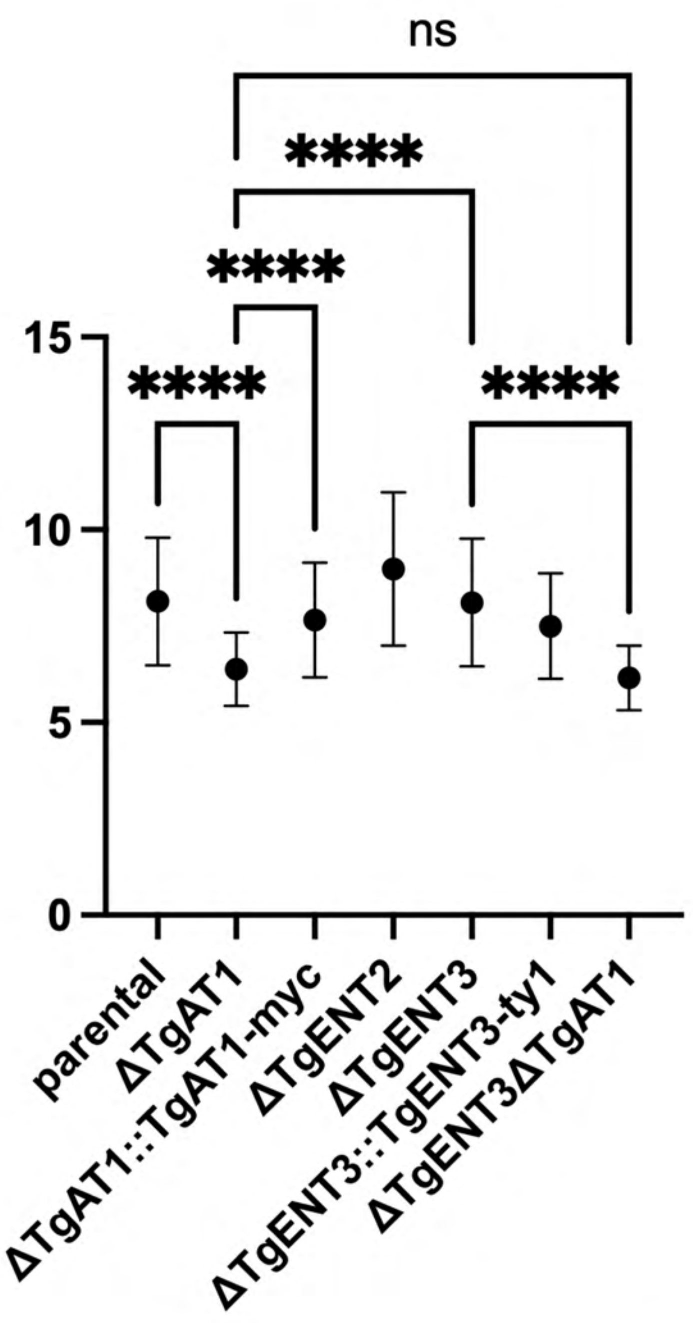

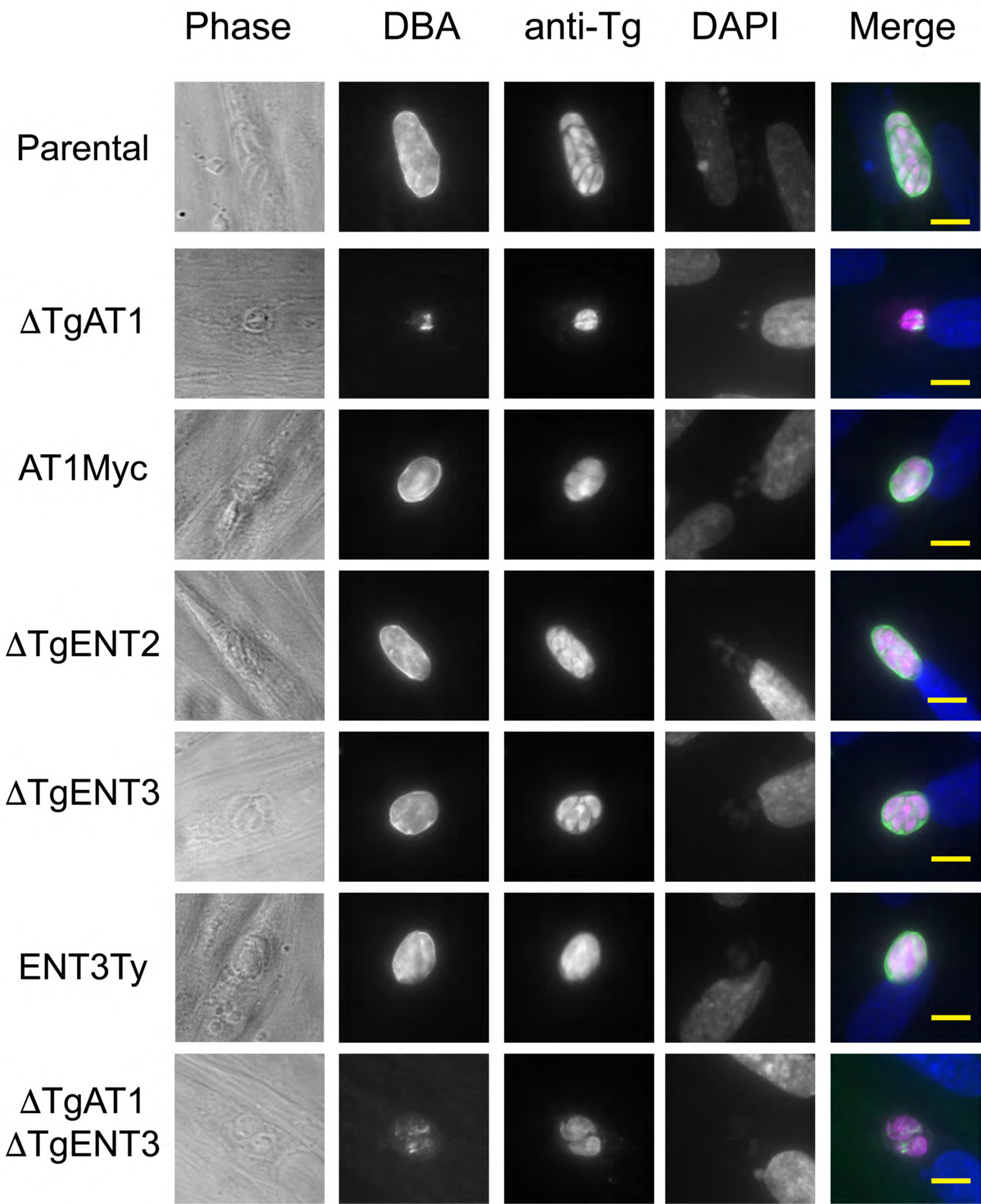

### Supplemental Figure 4

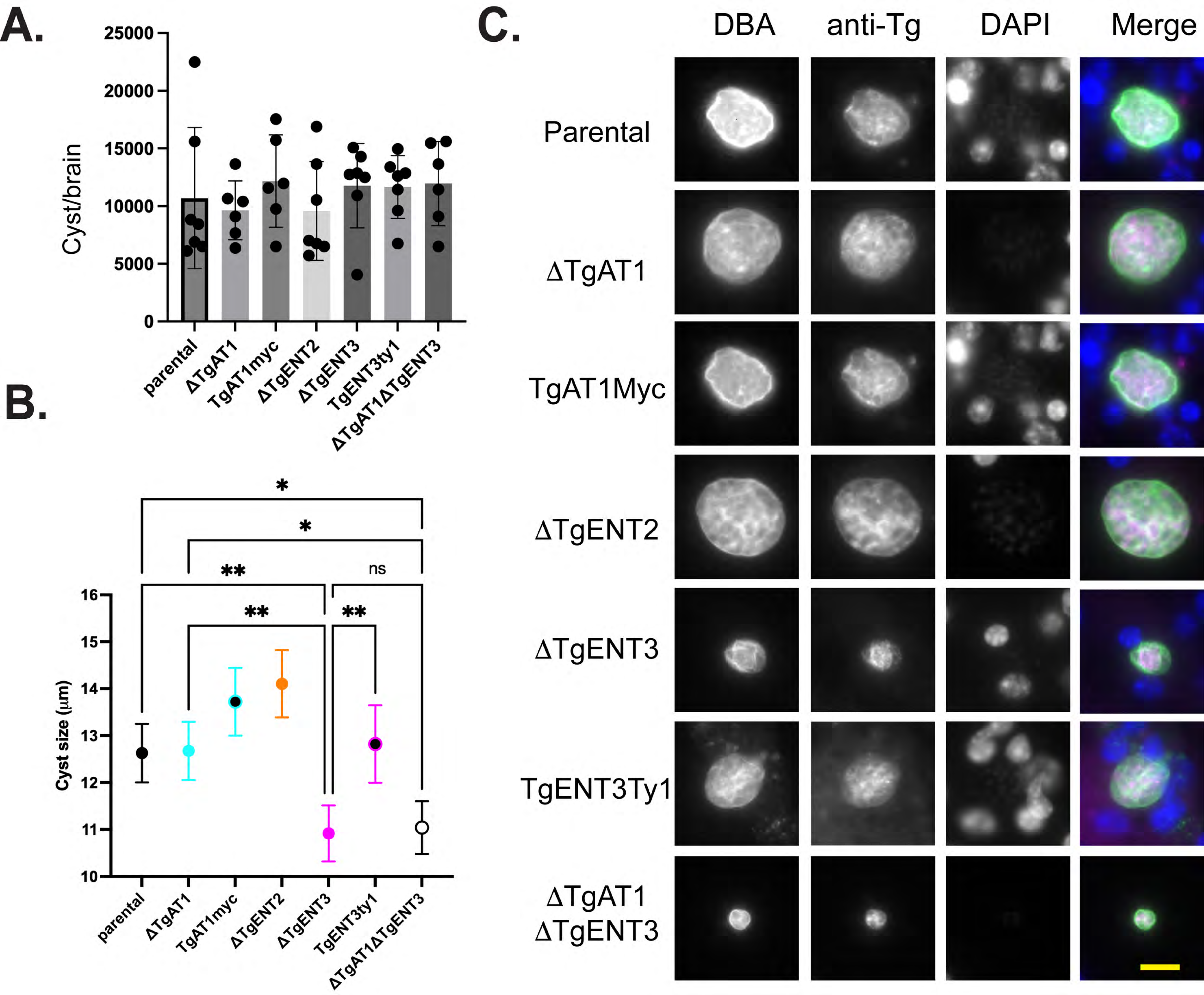
