## Supplemental Figure 5 for "Impact of Equilibrative Nucleoside Transporters on *Toxoplasma gondii* Infection and Differentiation"

A.

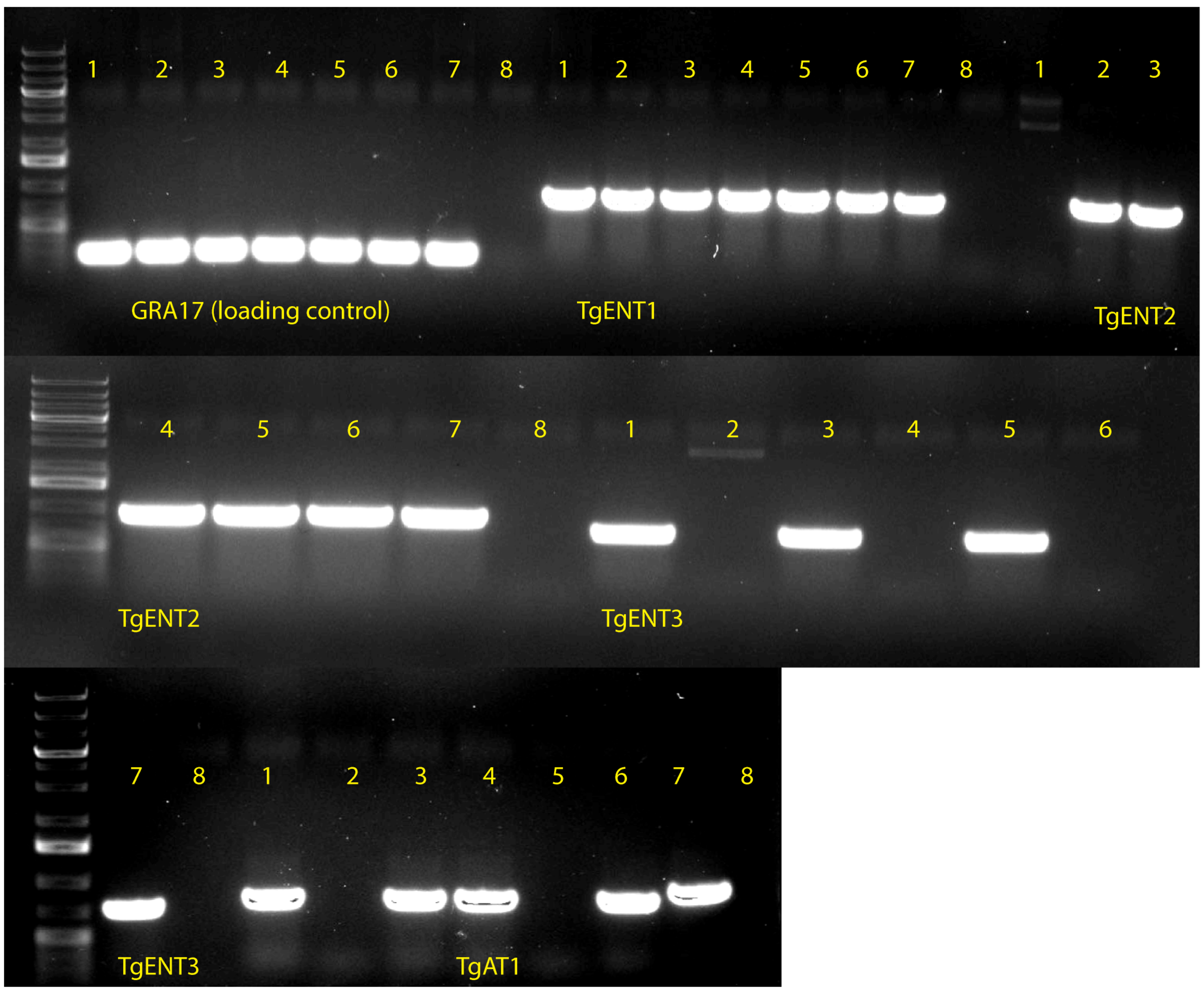


Each primer pair region has the following order of DNA templates
1. dENT2
2. dAT1dTgENT3
3. Parental
4. dENT3
5. dAT1
6. ENT3ty
7. AT1myc
8. Water control

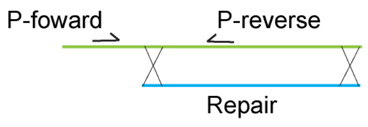


B.

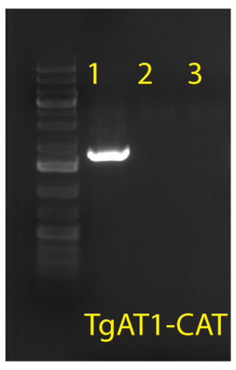


Primers targeting the chloamphenicol repair on AT1 KO parasites. Samples are:
1. dTgAT1
2. Parental ME49 strain
3. Water control

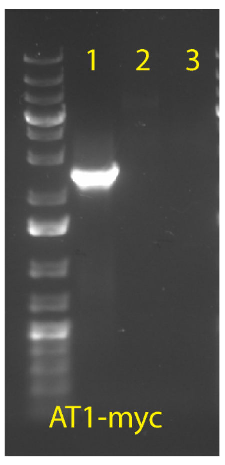


Primers targeting the AT1-myc repair on the complement AT1-myc parasites. Samples are:
1. AT1-myc
2. dTgAT1
3. Water control

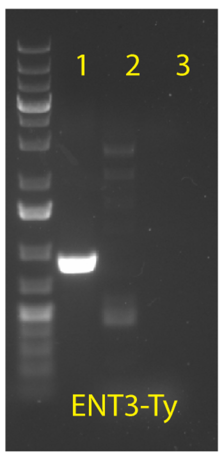


Primers targeting the ENT3-ty repair on the complement ENT3-ty parasites. Samples are:
1. ENT3-ty
2. dTgENT3
3. Water control

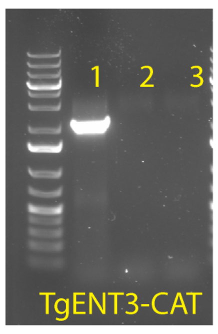


Primers targeting the chloamphenicol repair on ENT3 KO parasites. Samples are:
1. dTgENT3
2. Parental ME49 strain
3. Water control

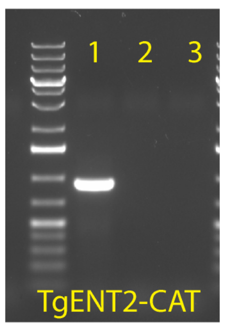


Primers targeting the chloamphenicol repair on ENT2 KO parasites. Samples are:
1. dTgENT2
2. Parental ME49 strain
3. Water control

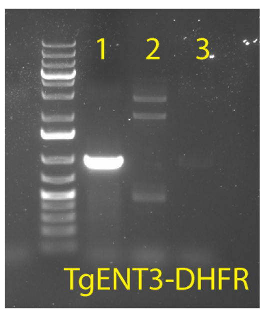


Primers targeting the DHFR repair on dAT1 dENT3 KO parasites. Samples are:
1. dAT1dENT3
2. dAT1
3. Water control

C.

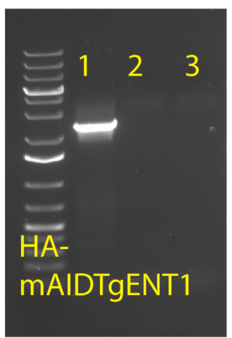


Each primer pair region has the following order of DNA templates
1. TgENT1-mAID-HA
2. Parental RH strain
3. Water control
